# Hepatic NF-kB-inducing Kinase (NIK) Suppresses Liver Regeneration in Chronic Liver Disease

**DOI:** 10.1101/238717

**Authors:** Yi Xiong, Adriana Souza Torsoni, Feihua Wu, Hong Shen, Yan Liu, Mark J Canet, M. Yatrik Shah, M. Bishr Omary, Yong Liu, Liangyou Rui

## Abstract

Hepatocyte replication maintains liver homeostasis and integrity. It is impaired in chronic liver disease, promoting disease progression. Herein, we have identified NF-kB-inducing kinase (NIK) as an unrecognized suppressor of hepatocyte replication. Hepatic NIK was aberrantly activated in chronic liver disease. Hepatocyte-specific deletion of *NIK* or its downstream mediator *IKKα* substantially accelerated hepatocyte proliferation and liver regeneration following partial hepatectomy. Mechanistically, NIK and IKKα suppressed the mitogenic JAK2/STAT3 pathway, thereby inhibiting hepatocyte cell cycle progression. Remarkably, inactivation of hepatic NIK largely reversed suppression of the hepatic JAK2/STAT3 pathway, hepatocyte replication, and liver regeneration induced by either chronic liver injury or metabolic stress. Our data suggest that hepatic NIK acts as a rheostat for liver regeneration to restrain liver overgrowth. Pathologic activation of hepatic NIK blocks hepatocyte replication, likely contributing to liver disease progression.

## Introduction

The liver is an essential metabolic organ, and often experiences metabolic stress during fasting and feeding and in overnutrition states (Rui, 2014). It is also frequently exposed to harmful insults, because it detoxifies endogenous and exogenous hepatotoxic substances. Dietary hepatotoxins and gut microbiota-derived toxic substances are transported directly to the liver through enterohepatic circulation, further increasing risk for liver injury. To compensate for a loss of hepatocytes, the liver has a powerful regenerative capability to maintain its homeostasis and integrity (Michalopoulos, 2017). After 70% of partial hepatectomy (PHx), rodents are able to regain their normal liver mass within one week (Miyaoka et al., 2012). Notably, it is equally important to avoid generation of aberrant liver cells from damaged hepatocytes in order to maintain liver integrity. Thus, a quality control mechanism likely exists to block injured hepatocytes from proliferating. We speculate that hepatocellular stress and/or injury signals activate hepatocyte-intrinsic sensors that in turn block proliferation of damaged hepatocytes through this putative quality control system.

Reparative hepatocyte proliferation is severely impaired in chronic liver disease, including nonalcoholic fatty liver disease (NAFLD), alcoholic liver disease, and a chronic exposure to hepatotoxins (Inaba et al., 2015; Michalopoulos, 2013; Richardson et al., 2007; Sancho-Bru et al., 2012). Hepatocyte proliferative arrest is associated with liver inflammation, injury, and fibrosis in patients with steatohepatitis (NASH) (Richardson et al., 2007), suggesting that impaired hepatocyte replication exacerbates disease progression. Numerous factors have been identified to promote hepatocyte proliferation, including various cytokines, growth factors, and the JAK2/STAT3, MAPK, PI 3-kinase, and NF-kB pathways (Michalopoulos, 2017). Paradoxically, most of these positive regulators are elevated in chronic liver disease. We reason that liver regeneration is also governed by negative regulators that function as a molecular rheostat to restrain liver overgrowth by counterbalancing positive regulators. Some of these negative regulators likely have dual functions and are involved in the quality control of liver regeneration by blocking proliferation of damaged hepatocytes. We postulate that in chronic liver disease, such negative regulators are overactivated by hepatocellular stress/injury, leading to pathological suppression of hepatocyte proliferation/liver regeneration. However, negative regulators for hepatocyte replication, in contrast to extensively studied positive regulators, are poorly understood.

In search for such negative regulators that couple hepatic injury to hepatocyte replication, we identified NF-kB-inducing kinase (NIK). NIK is a Ser/Thr kinase known to activate the noncanonical NF-kB2 pathway (Sun, 2012). It phosphorylates and activates IKKα (Xiao et al., 2001). IKKα phosphorylates NF-kB2 precursor p100, resulting in generation of a mature NF-kB2 p52 form (Sun, 2012; Xiao et al., 2001). We reported that metabolic stress, oxidative stress, hepatotoxins, and many cytokines all stimulate hepatic NIK (Jiang et al., 2015; Sheng et al., 2012). Consistently, hepatic NIK is aberrantly activated in both mice and humans with NAFLD, alcoholic liver disease, or other types of chronic liver disease (Shen et al., 2014). Therefore, NIK is involved in sensing of hepatocellular stress and damage, likely functioning as a hepatocyte-intrinsic sensor for stress/injury. In this work, we characterized hepatocyte-specific *NIK* (*NIK^Δhep^*) and *IKKα* (*IKKα^Δhep^*) knockout mice, and examined reparative hepatocyte replication using a PHx model. We found that the hepatic NIK/IKKα pathway suppresses reparative hepatocyte proliferation and liver regeneration by inhibiting the JAK2/STAT3 pathway. Aberrantly activated hepatic NIK in chronic liver disease is responsible for, in part, impairment in liver regeneration. Our data suggest that NIK is an unrecognized hepatocyte-intrinsic sensor for stress/injury and negative regulator of hepatocyte proliferation.

## Results

### Hepatocyte-specific deletion of *NIK* accelerates liver regeneration in mice

To assess the role of hepatic NIK in hepatocyte reparative proliferation, we performed 70% of PHx on hepatocyte-specific *NIK* knockout mice at age of 8 weeks (Mitchell and Willenbring, 2008). *NIK^Δhep^* mice were generated by crossing *NIK^flox/flox^* mice with *albumin-Cre* drivers as described previously (Shen et al., 2017). Proliferating cells were detected by immunostaining liver sections with antibody against Ki67, a marker of proliferating cells (Fig. 1A). Liver proliferating rates were low (<1%) under basal conditions and comparable between *NIK^Δhep^* and *NIK^flox//flox^* mice (Fig. 1B). PHx markedly increased the number of Ki67^+^ proliferating cells in both groups 48 h after PHx; remarkably, liver proliferating cells were 85% higher in *NIK^Δhep^* mice relative to *NIK^flox/flox^* mice (Fig. 1B). In line with these findings, the number of liver BrdU+ proliferating cell, as assessed by BrdU assays, was also much higher in *NIK^Δhep^* than in *NIK^flox/flox^* mice (Fig. 1C). Liver cell proliferation declined 48 h after PHx in both *NIK^Δhep^* and *NIK^flox/flox^* mice, and became comparable between these two groups within 96 h after PHx (Fig. 1B).

**Figure 1.**
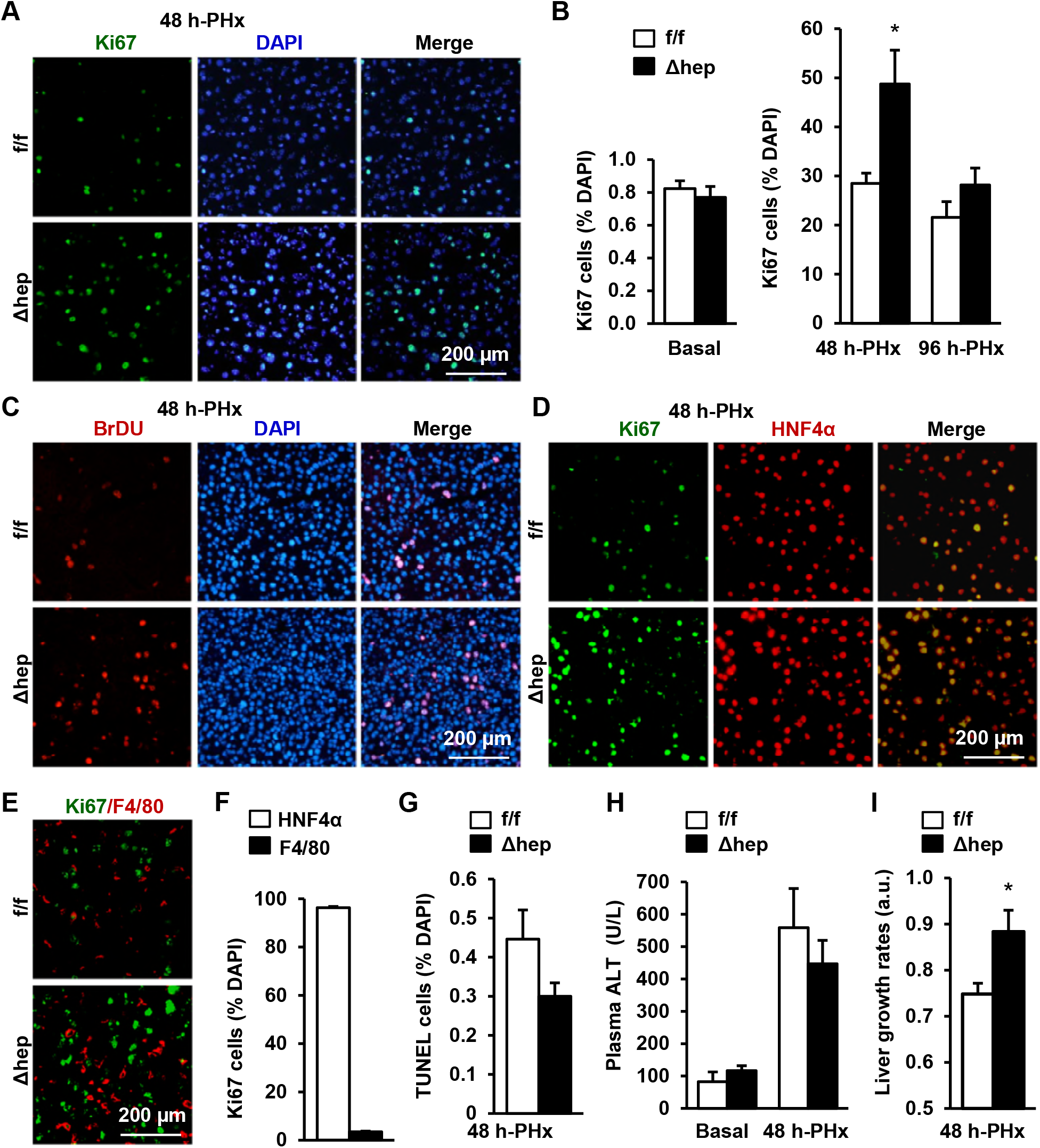
Hepatocyte-specific deletion of *NIK* accelerates hepatocyte reparative proliferation. *NIK^flox/flox^*(n=7) and *NIK^Δhep^* (n=7) male mice (8 weeks) were subjected to PHx, and livers were harvested 48 h or 96 h later. (**A**) Representative immunostaining of liver sections (48 h after PHx) with anti-Ki67. (**B**) Ki67^+^ cells were counted and normalized to total DAPI+ cells. (C) Representative immunostaining of liver sections (48 h after PHx) with anti-BrdU antibodies. (**D-E**) Representative images of liver sections (48 h after PHx) costained with anti-Ki67 and anti-HNF4α antibodies (**D**) or anti-Ki67 and anti-F4/80 antibodies (**E**). (**F**) Ki67^+^HNF4α+ and Ki67^+^F4/80+ cells were counted and normalized to total Ki67^+^ cells. (**G**) Liver cell death were assessed 48 h after PHx using TUNEL reagents. (**H**) Plasma ALT levels. (**I**) Liver growth rates within 48 h after PHx. Data were statistically analyzed with two-tailed Student’s t test, and presented as mean ± SEM. *p<0.05.

To confirm that the proliferating cells are hepatocytes, we costained liver sections with antibodies against either Ki67 and HNF4α (a hepatocyte marker) or Ki67 and F4/80 (a Kupffer cell/macrophage marker). HNF4α^+^ hepatocytes accounted for 96% of liver Ki67^+^ proliferating cells in *NIK^Δhep^* mice 48 h after PHx (Figs. 1D and 1F), whereas F4/80^+^ Kupffer cells/macrophages accounted for <4% of Ki67^+^ cells (Fig. 1E-F). Together, these data indicate that NIK is an intrinsic suppressor of hepatocyte proliferation.

Next, we examined the effect of NIK deficiency on hepatocyte death. The number of liver TUNEL^+^ apoptotic cells, as assessed by TUNEL assays, was slightly lower in *NIK^Δhep^* than in *NIK^flox/flox^* mice, but the difference was not statistically significant (Fig. 1G). Plasma alanine aminotransferase (ALT) activity, a liver injury index, was comparable between *NIK^Δhep^* and *NIK^flox//flox^* mice under both basal and PHx conditions (Fig. 1H). Thus, it is unlikely that hepatic NIK regulates hepatocyte death under these conditions.

To further validate the role of hepatic NIK in the maintenance of liver homeostasis, we quantified liver regeneration rates within 48 h after PHx. In line with increased hepatocyte proliferation, liver growth rates were also significantly increased in *NIK^Δhep^* mice compared with *NIK^flox/flox^* mice (Fig. 1I). In light of these findings, we propose that NIK is a hepatocyte-intrinsic rheostat for reparative proliferation that is involved in the maintenance of liver homeostasis and integrity. Moreover, damage-induced NIK activation likely provides a quality control mechanism to prevent generation of aberrant cells from damaged hepatocytes.

### The NF-kB1, MAPK, and PI 3-kinase pathways do not mediate NIK suppression of hepatocyte reparative proliferation

To further confirm NIK inhibition of hepatic cell cycle progression, we measured the levels of hepatic cyclin D1, which is believed to drive hepatocyte proliferation after PHx (Michalopoulos, 2013). Hepatic cyclin D1 was undetectable under basal conditions in both *NIK^Δhep^* and *NIK^flox/flox^* mice, and markedly increased after PHx in both groups (Fig. 2A). In line with increased hepatocyte proliferation, hepatic cyclin D1 levels were significantly higher in *NIK^Δhep^* than in *NIK^flox/flox^* mice (Fig. 2A-B).

**Figure 2.**
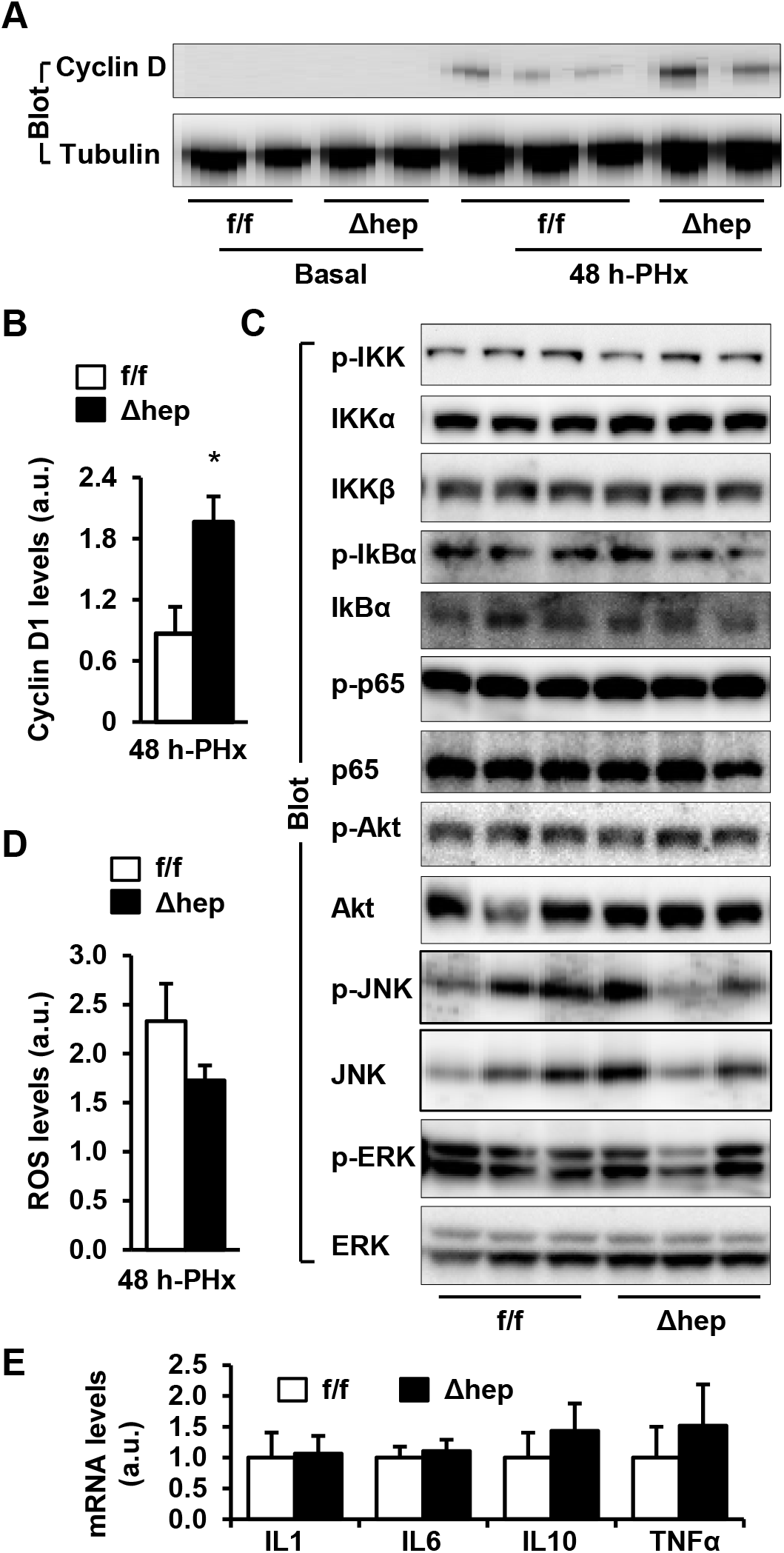
Hepatic NIK deficiency upregulates cyclin D1 without altering NF-kB1, Akt, and MAPK pathways in the liver. *NIK*^flox/flox^ and *NIK^Δhep^* male mice (8 weeks) were subjected to PHx. (**A-B**) Liver extracts were immunoblotted with anti-cyclin D1 antibody (48 h after PHx). Cyclin D1 levels were quantified and normalized to α-tubulin levels (*NIK^flox/flox^*: n=7, *NIK^Δhep^*: n=7). (**C**) Liver extracts were immunoblotted with the indicated antibodies (4 h after PHx). (**D**) Liver ROS levels 48 h after PHx (normalized to liver weight). *NIK^flox/flox^*: n=4, *NIK^Δhep^*: n=4. (**E**) Liver cytokine expression was measured by qPCR and normalized to 36B4 expression (48 h after PHx). *NIK^flox/flox^*: n=5, *NIK^Δhep^*: n=5. Data were statistically analyzed with two-tailed Student’s t test, and presented as mean ± SEM. *p<0.05.

We next sought to study the molecular mechanism of NIK action. NF-kB1, MAPK, and PI 3-kinase pathways are known to be involved in mediating PHx-stimulated liver regeneration (Michalopoulos, 2013; Pauta et al., 2016; Wuestefeld et al., 2013). Surprisingly, phosphorylation of hepatic IKKα/β, IkBα, p65 (the NF-kB1 pathway), Akt (pSer473) (the PI 3-kinase pathway), ERK1/2, and JNK (the MAPK pathway) was comparable between *NIK^Δhep^* and *NIK^flox/flox^* mice 4 h after PHx (Fig. 2C). We also assessed liver reactive oxygen species (ROS) and expression of various cytokines, and did not detect difference between *NIK^Δhep^* and *NIK^flox/flox^* mice (Fig. 2D-E). Therefore, these pathways are unlikely to mediate suppression of liver regeneration by hepatic NIK.

### NIK directly suppresses the Janus kinase 2 (JAK2)/STAT3 pathway

JAK2 phosphorylates and activates STAT3, and the JAK2/STAT3 pathway is believed to drive hepatocyte proliferation (Shi et al., 2017; Wang et al., 2011). We postulated that NIK might suppresses hepatocyte proliferation by inhibiting the JAK2/STAT3 pathway. Liver extracts were prepared 4 h after PHx and immunoblotted with anti-phospho-JAK2 (pTyr1007/1008) or anti-phospho-STAT3 (pTyr705) antibodies. Phosphorylation of both JAK2 and STAT3 was significantly higher in *NIK^Δhep^* than in *NIK^flox/flox^* littermates (Fig. 3A).

**Figure 3.**
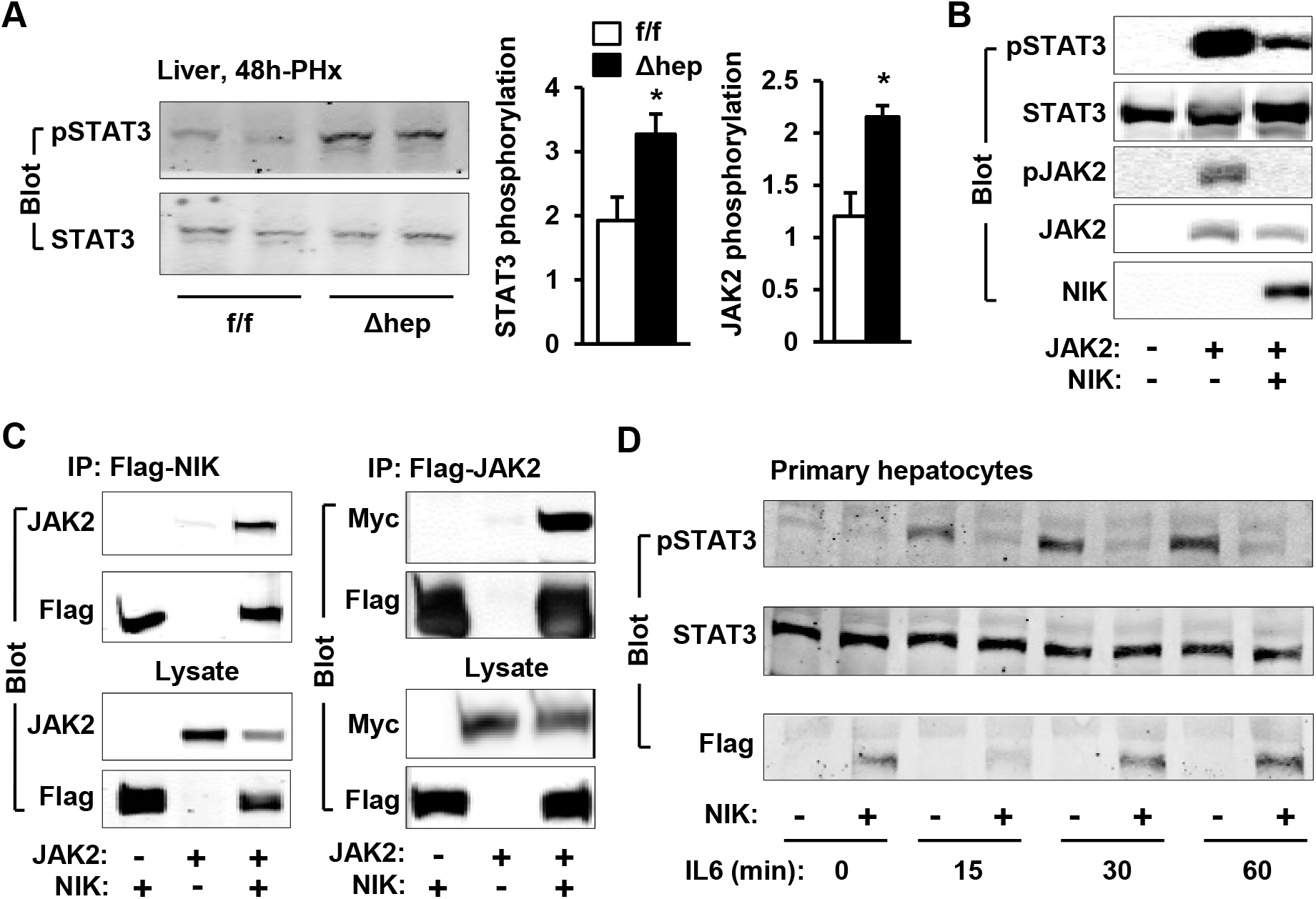
NIK inhibits the JAK2/STAT3 pathway. (**A**) Liver extracts were prepared from *NIK^flox/flox^* and *NIK^Δhep^* male 4 h after PHx and immunoblotted with anti-phospho-JAK2 and anti-phospho-STAT3 antibodies. Phosphorylation of JAK2 (pTyr1007/1008) and STAT3 (pTyr705) was normalized to total JAK2 and STAT3 levels, respectively. (**B**) STAT3 and JAK2 were coexpressed with or without NIK in HEK293 cells. Cell extracts were immunoblotted with the indicated antibodies. (**C**) NIK was coexpressed with JAK2 in HEK293 cells. Cell extracts were immunoprecipitated (IP) and immunoblotted with the indicated antibodies. (**D**) Mouse primary hepatocytes were transduced with NIK or β-gal adenoviral vectors and stimulated with IL6 (10 ng/ml). Cell extracts were immunoblotted with the indicated antibodies. Data were statistically analyzed with two-tailed Student’s t test, and presented as mean ± SEM. *p<0.05.

To confirm that NIK directly inhibits the JAK2/STAT3 pathway, we transiently coexpressed JAK2 and STAT3 with NIK in HEK293 cells. JAK2 was autophosphorylated and robustly phosphorylated STAT3 in the absence of NIK (Fig. 3B), as we previously reported (Rui and Carter-Su, 1999). Overexpression of NIK dramatically attenuated phosphorylation of both JAK2 and STAT3 (Fig. 3B). Moreover, NIK was coimmunoprecipitated with JAK2 (Fig. 3C). These data indicate that NIK binds to JAK2 and inhibits JAK2 activity, thereby suppressing the JAK2/STAT3 pathway.

We next tested if NIK negatively regulates interleukin 6 (IL6)-stimulated activation of the JAK2/STAT3 pathway, because the IL6/JAK2/STAT3 cascade is required for hepatocyte reparative proliferation (Cressman et al., 1996; Riehle et al., 2008). Mouse primary hepatocytes were transduced with NIK or β-galactosidase (β-gal; control) adenoviral vectors, and then stimulated with IL6. IL6 rapidly and robustly stimulated phosphorylation of STAT3 in β-gal-transduced hepatocytes; strikingly, overexpression of NIK completely blocked IL6-stimulated phosphorylation of STAT3 (Fig. 3D). We did not detect endogenous JAK2, because its levels were below the detection threshold of our assays. Overall, our data unveiled unrecognized crosstalk between NIK and JAK2/STAT3 pathways. NIK inhibits hepatocyte proliferation at least in part by restraining the JAK2/STAT3 pathway.

### Hepatic IKKα suppresses liver regeneration after PHx

NIK phosphorylates and activates IKKα (Sun, 2012), prompting us to test if hepatocyte-specific *IKKα* knockout mice, like *NIK^Δhep^* mice, also display accelerated liver regeneration. *IKKα^Δhep^* mice were generated by crossing *IKKα^flox/flox^* mice with *albumin-Cre* drivers (Liu et al., 2008). *IKKα* was disrupted specifically in the liver, but not the brain, heart, kidney, skeletal muscle, and spleen, of *IKKα^Δhep^* mice (Fig. 4A). We performed PHx on *IKKα^flox/flox^* and *IKKα^Δhep^* male mice at 8-9 weeks of age. The number of liver Ki67^+^ proliferating cell was significantly higher in *IKKα^Δhep^* than in *IKKα^flox/flox^* littermates 48 h after PHx (Fig. 4B). The proliferating cells were HNF4α+ hepatocytes (Fig. 4C). Consistently, liver cyclin D1 levels were significantly higher in *IKKα^Δhep^* than in *IKKα^flox/flox^* mice (Fig. 4D). In contrast, liver cell death was comparable between *IKKα^Δhep^* and *IKKα^flox/flox^* littermates (Fig. 4E). Consequently, liver regeneration rates were significantly higher in *IKKα^Δhep^* mice relative to *IKKα^flox/flox^* mice (Fig. 4F). These results suggest that IKKα acts downstream of NIK to suppress liver regeneration.

**Figure 4.**
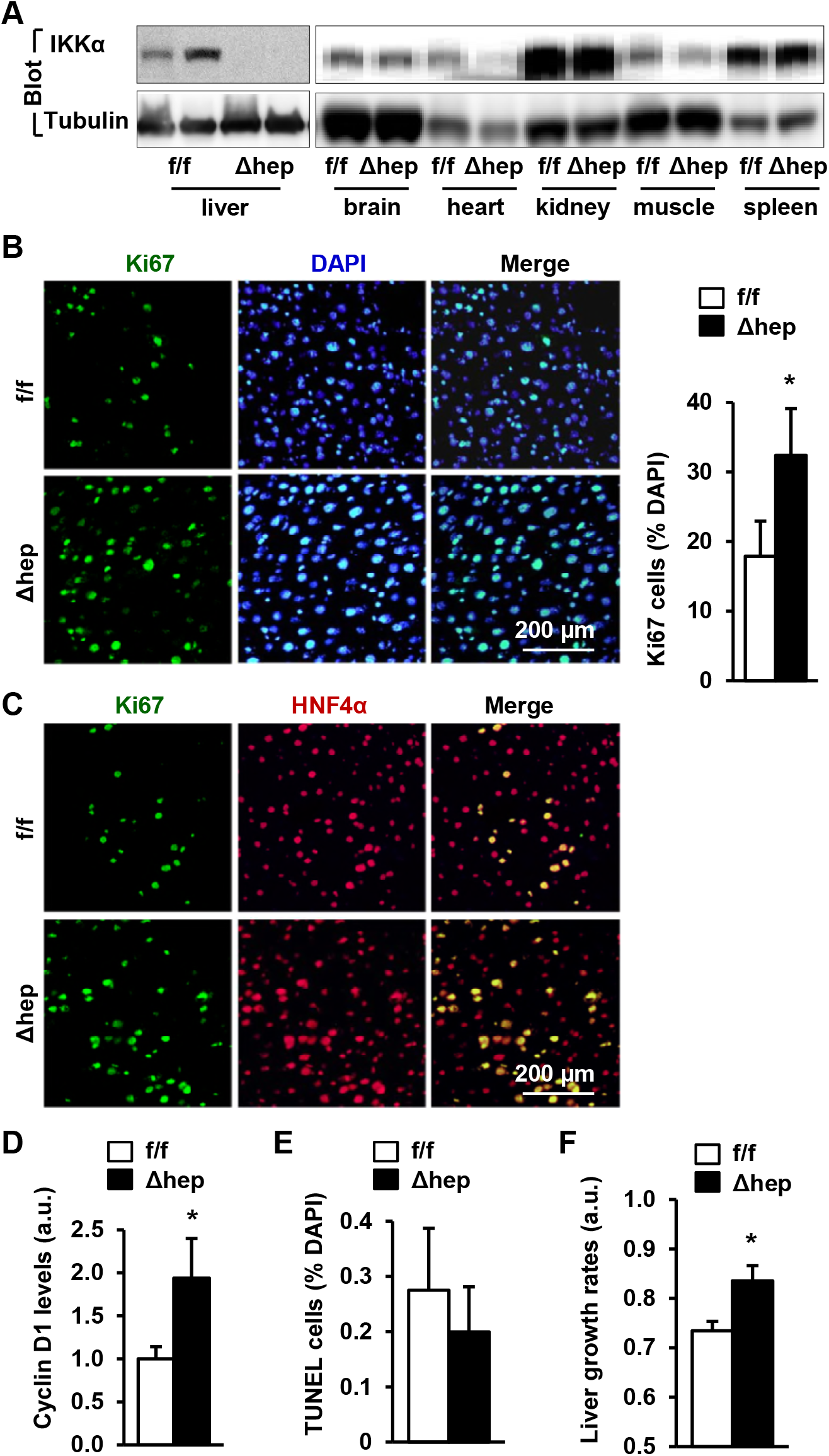
Hepatocyte-specific deletion of *IKKα* accelerates hepatocyte reparative proliferation. (**A**) Tissue extracts were immunoblotted with anti-IKKα or anti-α-tubulin antibodies. (**B-F**) *IKKα^flox/flox^* (n=6) and *IKKα^Δhep^* (n=6) male littermates were subjected to PHx, and livers were harvested 48 h later. (**B**) Liver sections were immunostained with anti-Ki67 antibody, and Ki67^+^ cells were counted and normalized to total DAPI^+^ cells. (**C**) Representative images of liver sections costained with anti-Ki67 and anti-HNF4α antibodies. (**D**) Liver cyclin D1 was measured by immunoblotting (normalized to α-tubulin levels). (**E**) TUNEL-positive cells in liver sections. (**F**) Liver growth rates within 48 h after PHx. Data were statistically analyzed with two-tailed Student’s t test, and presented as mean ± SEM. *p<0.05.

We next sought to test if IKKα, like NIK, inhibits the JAK2/STAT3 pathway. Phosphorylation of both JAK2 and STAT3 was significantly higher in *IKKα^Δhep^* than in *IKKα^flox/flox^* mice 4 h after PHx (Fig. 5A-B). To determine whether IKKα directly inhibits JAK2, IKKα was transiently coexpressed with JAK2 in HEK293 cells. IKKα markedly decreased JAK2 autophosphorylation and the ability of JAK2 to phosphorylate STAT3 (Fig. 5C). Furthermore, IKKα was coimmunoprecipitated with JAK2 (Fig. 5D). Thus, NIK is able to inhibit the JAK2/STAT3 pathway both directly and indirectly through activating IKKα.

**Figure 5.**
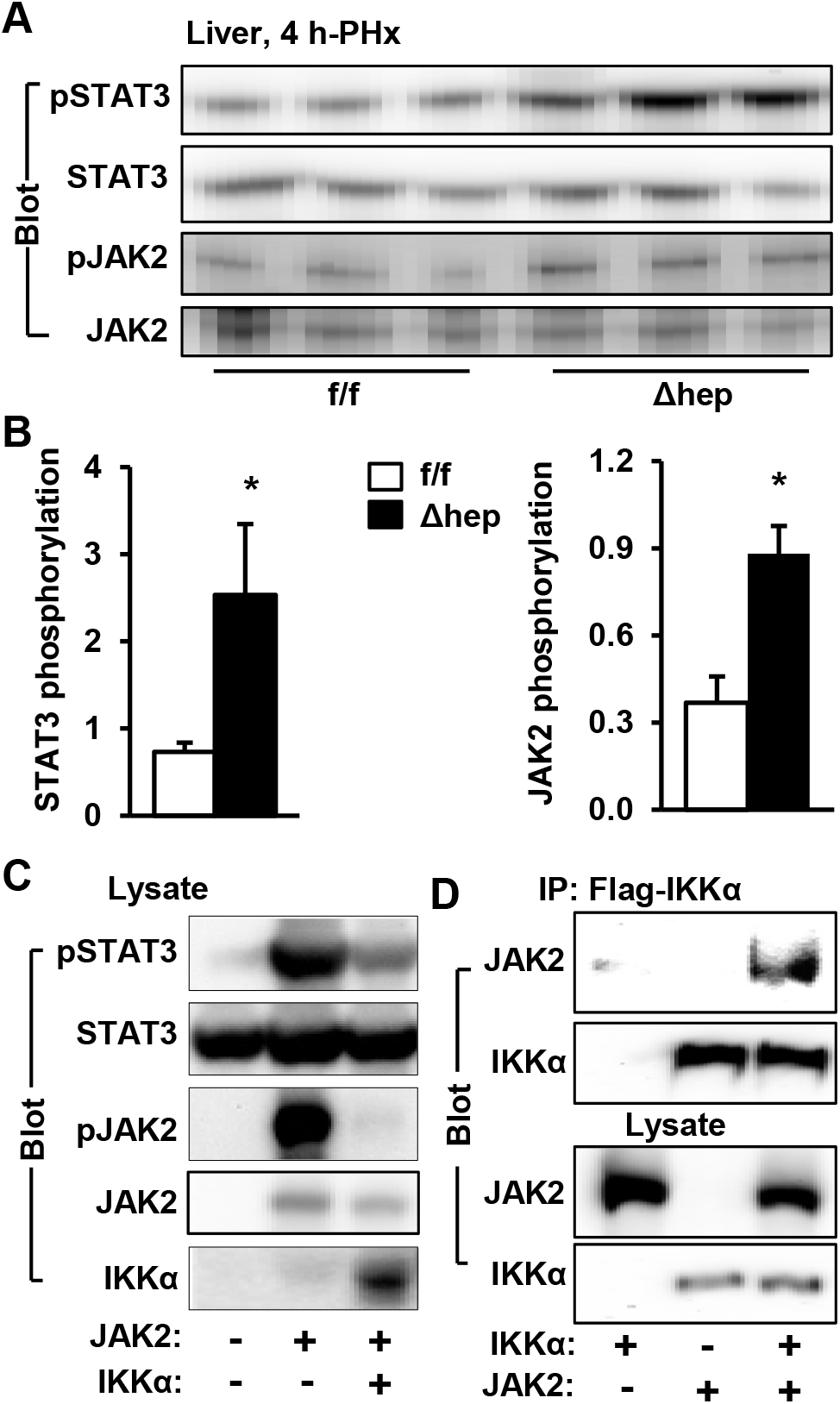
IKKα inhibits the JAK2/STAT3 pathway. (**A-B**) Liver extracts were prepared 4 h after PHx and immunoblotted with anti-phospho-JAK2 and anti-phospho-STAT3 antibodies. Phosphorylation of JAK2 (pTyr1007/1008) and STAT3 (pTyr705) was normalized to total JAK2 and STAT3 levels, respectively. *IKKα^flox/flox^*: n=5, *IKKα^Δhep^*: n=5. (**C**) STAT3 and JAK2 were coexpressed with IKKα in HEK293 cells. Cell extracts were immunoblotted with the indicated antibodies. (**D**) IKKα and JAK2 were coexpressed in HEK293 cells. Cell extracts were immunoprecipitated (IP) and immunoblotted with the indicated antibodies. Data were statistically analyzed with two-tailed Student’s t test, and presented as mean ± SEM. *p<0.05.

### Deletion of hepatic *NIK* reverses hepatotoxin-induced suppression of liver regeneration

Since hepatic NIK is aberrantly activated in chronic liver disease (Shen et al., 2014; Sheng et al., 2012), we speculated that it might be a causal factor for impaired liver regeneration that promotes disease progression. To model chronic liver disease, mice were treated with 2-acetylaminofluorene (AAF), a hepatotoxin (Laishes and Rolfe, 1981). AAF treatment considerably activated hepatic NIK, as assessed by upregulation of NF-kB2 p52 (Fig. 6A). To examine the impact of elevated NIK on liver regeneration, *NIK^Δhep^* and *NIK^flox/flox^* mice were pretreated with AAF for 10 days prior to PHx. Proliferating cells were assessed 48 h after PHx by immunostaining liver sections with anti-Ki67 antibody (Fig. 6B). Baseline hepatocyte proliferation was low and similar between *NIK^flox/flox^* and *NIK^Δhep^* mice prior to PHx (Fig. 6C). PHx markedly increased hepatocyte proliferation in PBS-treated *NIK^flox/flox^* mice (control) as expected, and AAF pretreatment substantially decreased hepatocyte proliferation by >50% (Fig. 6D). Remarkably, deletion of hepatic *NIK* largely reversed AAF-induced suppression of hepatocyte proliferation in *NIK^Δhep^* mice (Fig. 6D). In contrast, plasma ALT levels (a liver injury index) was similar between *NIK^flox/flox^* and *NIK^Δhep^* mice (Fig. 6E). In line with increased hepatocyte proliferation, liver regeneration rates after PHx were also significantly higher in AAF-pretreated *NIK^Δhep^* mice relative to *NIK^flox/flox^* littermates (Fig. 6F).

**Figure 6.**
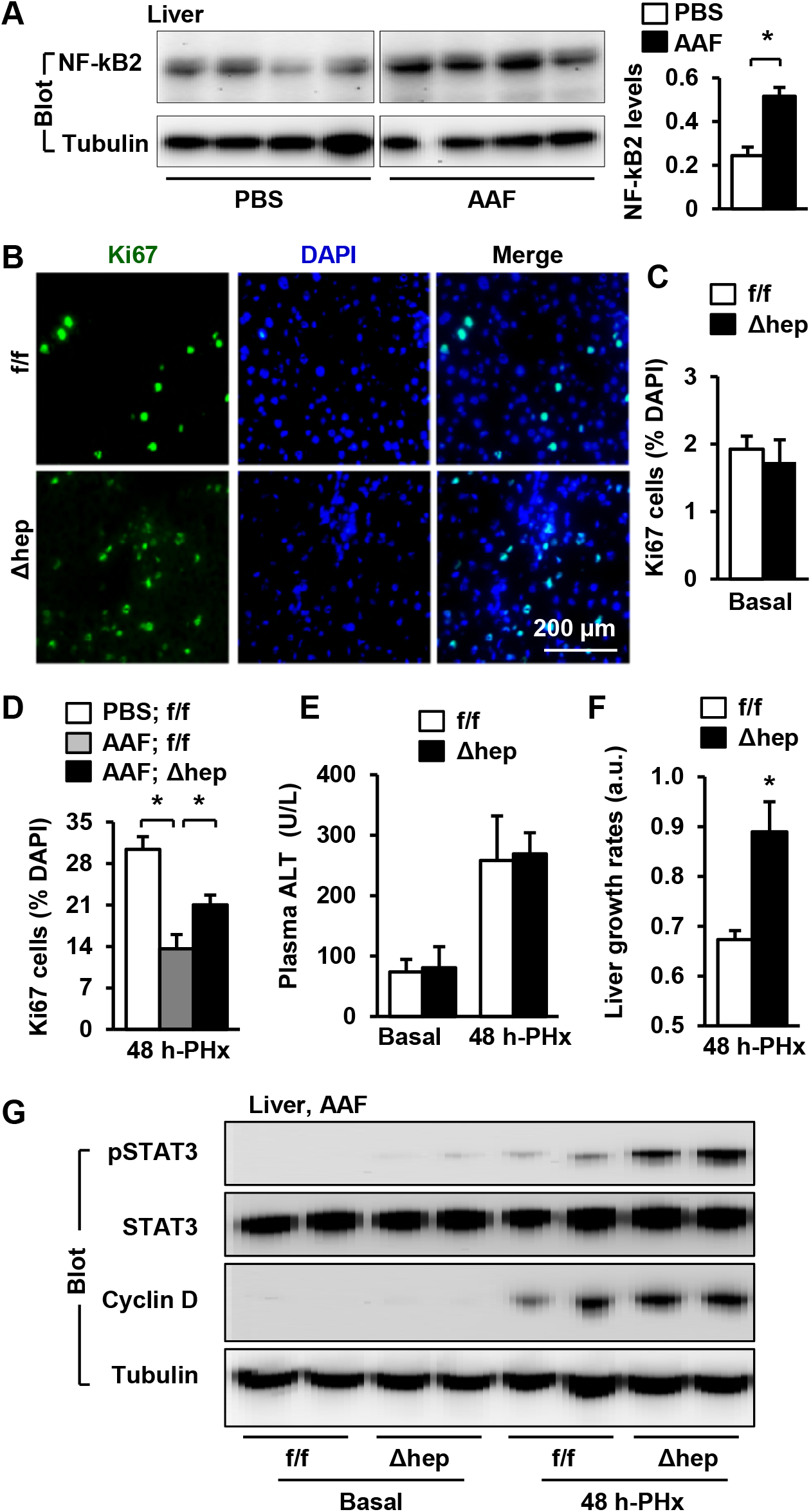
Hepatocyte-specific deletion of *NIK* reverses AAF-induced impairment in hepatocyte reparative proliferation. (**A**) C57BL/6 males (8 weeks) were treated with PBS or AAF (10 mg/kg body weight, gavage) daily for 10 days. NF-kB2 p52 in liver extracts was immunoblotted with anti-NF-kB2 antibody (normalized to α-tubulin levels). PBS: n=4, AAF: n=4. (**B-G**) *NIK^flox/flox^* and *NIK^Δhep^* males were treated with PBS or AAF (10 mg/kg body weight) for 10 days and then subjected to PHx. Livers were harvested 48 h later. (**B**) Representative immunostaining of liver sections with anti-Ki67 antibody. (**C**) Baseline Ki67^+^ cell number in resected liver sections obtained from PHx. *NIK^flox/flox^*: n=3, *NIK^Δhep^:* n=3. (**D**) Ki67^+^ cell number in liver sections (normalized to DAPI+ cells). PBS;*NIK^flox/flox^*: n=3, AAF;*NIK^flox/flox^*: n=5, AAF;*NIK^Δhep^*: n=5. (**E**) Plasma ALT levels. *NIK^flox/flox^*: n=3, *NIK^Δhep^*: n=4. (**F**) Liver regeneration rates within 48 h after PHx. *NIK^flox/flox^*: n=5, *NIK^Δhep^*: n=5. (**G**) Liver extracts were immunoblotted with the indicated antibodies. Data were statistically analyzed with two-tailed Student’s t test, and presented as mean ± SEM. *p<0.05.

We next examined cell signaling events that drive cell cycle progression. We detected baseline phosphorylation of hepatic STAT3 in AAF-pretreated *NIK^Δhep^* but not *NIK^flox/flox^* mice prior to PHx (Fig. 6G). PHx stimulated phosphorylation of hepatic STAT3 in both *NIK^Δhep^* and *NIK^flox/flox^* mice; however, STAT3 phosphorylation was substantially higher in *NIK^Δhep^* mice (Fig. 6G). Baseline hepatic cyclin D was undetectable in both AAF-pretreated *NIK^Δhep^* and *NIK^flox/flox^* mice prior to PHx. PHx upregulated hepatic cyclin D1 to a higher level in *NIK^Δhep^* mice relative to *NIK^flox/flox^* littermates (Fig. 6G). Together, these data further support the notion that NIK serves as a hepatocyte-intrinsic rheostat to restrain liver regeneration through inhibiting the JAK2/STAT3 pathway. Importantly, abnormal activation of hepatic NIK is likely responsible for impaired liver regeneration in chronic liver disease, contributing to disease progression.

### Deletion of hepatic *NIK* reverses NAFLD-associated suppression of liver regeneration

To model NAFLD, *NIK^Δhep^* and *NIK^flox/flox^* mice were fed a high fat diet (HFD) for 10 weeks. HFD-fed *NIK^Δhep^* and *NIK^flox/flox^* mice developed liver steatosis, as assessed by liver triacylglycerol (TAG) levels, to a similar degree (Fig. 7A). HFD feeding was reported to increase hepatic NIK activity (Sheng et al., 2012). Consistently, NF-kB2 p52 levels was higher in HFD-fed than normal chow-fed mice (Fig. 7B).

**Figure 7.**
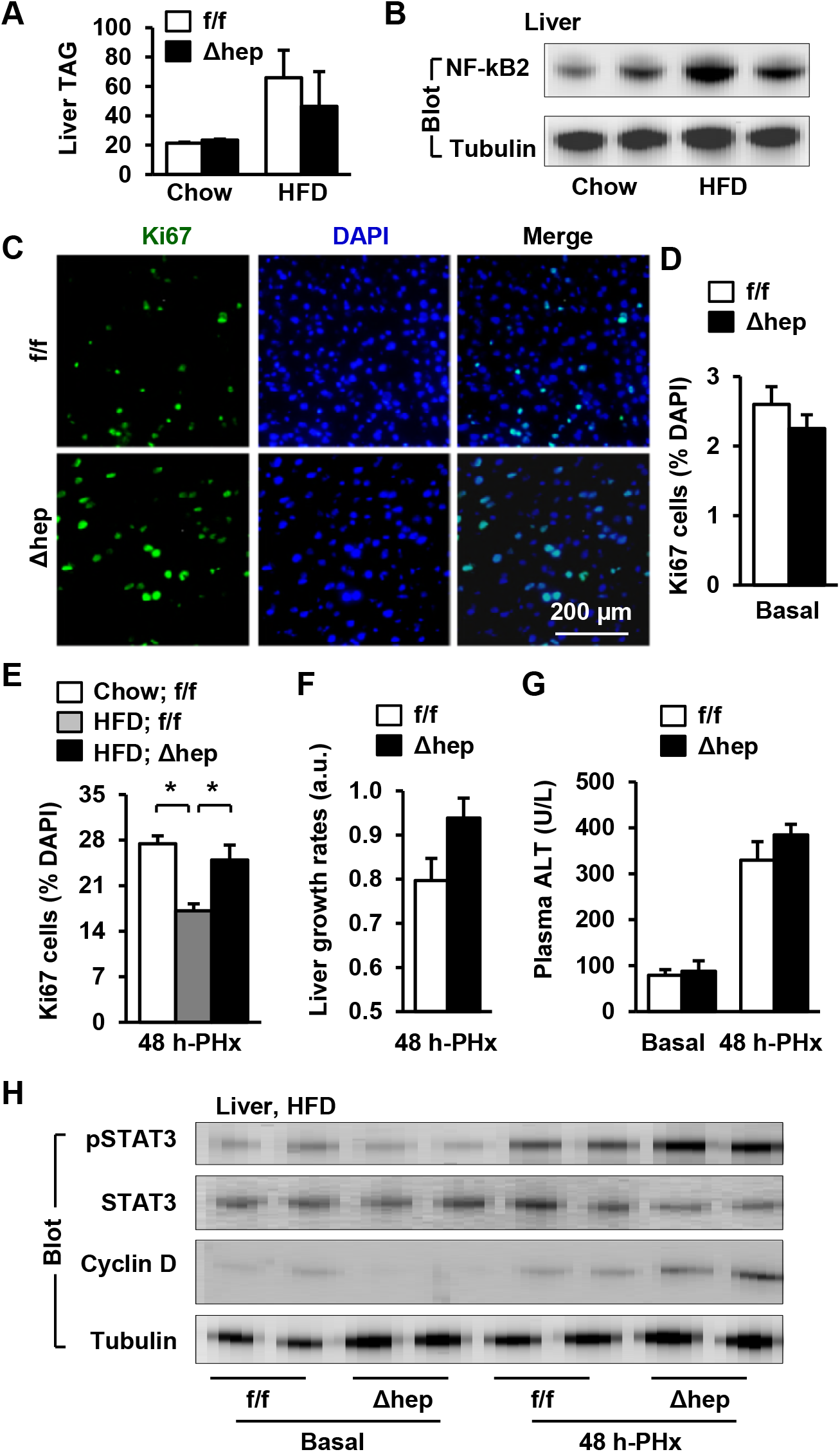
Hepatic NIK deficiency corrects impaired hepatocyte reparative proliferation in mice with NAFLD. (**A-B**) C57BL/6 males (8 weeks) were fed a normal chow diet (n=5) or a HFD (n=5) for 10 weeks. (**A**) Liver TAG levels (normalized to liver weight). (**B**) NF-kB2 p52 in liver extracts was immunoblotted with anti-NF-kB2 antibody (normalized to α-tubulin levels). (**C-H**) *NIK^flox/flox^* and *NIK^Δhep^* males were fed a HFD for 10 weeks followed by PHx, and livers were harvested 48 h after PHx. (**C**) Representative immunostaining of liver sections with anti-Ki67 antibody. (**D**) Baseline Ki67^+^ cell number in resected liver sections obtained from PHx. *NIK^flox/flox^*: n=4, *NIK^Δhep^*: n=4. (**E**) Liver Ki67^+^ cell number (normalized to DAPI+ cells). Chow;NIK^flox/flox^: n=3, HFD;NIK^flox/flox^: n=5, HFD;NIK^Δhep^: n=5. (**F**) Liver growth rates within 48 h after PHx. *NIK^flox/flox^*: n=4, *NIK^Δhep^*: n=5. (G) Plasma ALT levels. *NIK^flox/flox^*: n=3, *NIK^Δhep^*: n=4. (H) Liver extracts were immunoblotted with the indicated antibodies. Data were statistically analyzed with two-tailed Student’s t test, and presented as mean ± SEM. *p<0.05.

NAFLD/NASH is associated with impaired liver regeneration (Collin de l’Hortet et al., 2014; Inaba et al., 2015). We hypothesized that elevated hepatic NIK is responsible for suppression of liver generation in NAFLD/NASH. *NIK^Δhep^* and *NIK^flox/flox^* mice were fed a HFD for 10 weeks followed by PHx. Hepatocyte proliferation was assessed 48 h post-PHx by staining liver sections with anti-Ki67 antibody (Fig. 7C). Baseline hepatocyte proliferation was similarly low in both *NIK^Δhep^* and *NIK^flox/flox^* mice prior to PHx (Fig. 7D). PHx markedly induced hepatocyte proliferation in chow-fed *NIK^flox/flox^* mice, and HFD feeding substantially decreased Ki67^+^ proliferating hepatocyte number as expected (Fig. 7E). Remarkably, hepatocyte-specific deletion of *NIK* dramatically increased Ki67^+^ liver cell number in *NIK^Δhep^* mice close to normal levels (Fig. 7E). Consistently, liver growth rates was higher in *NIK^Δhep^* mice relative to *NIK^flox/flox^* mice, albeit those differences were not statistically significant (Fig. 7F). It is worth mentioning that high levels of liver TAG in HFD-fed mice might mask our assessments of liver growth rates that were based on liver weight changes. Plasma ALT levels were comparable between *NIK^Δhep^* and *NIK^flox/flox^* littermates under both basal and PHx conditions (Fig. 7G), suggesting that hepatic NIK does not directly affect liver injury under these conditions.

We further explored mitogenic pathways in the livers of these mice. STAT3 phosphorylation was similar between HFD-fed *NIK^Δhep^* and *NIK^flox/flox^* mice under baseline conditions, and increased to a considerably higher level in *NIK^Δhep^* mice relative to *NIK^flox/flox^* mice 48 h post-PHx (Fig. 7H). Hepatic cyclin D1 levels were also higher in *NIK^Δhep^* mice relative to *NIK^flox/flox^* mice after PHx (Fig. 7H). Overall, these data suggest that aberrant activation of hepatic NIK in NAFLD/NASH suppresses hepatocyte reparative proliferation through inhibiting the JAK2/STAT3 pathway.

## Discussion

Reparative hepatocyte proliferation supplies new hepatocytes to replace lost and damaged hepatocytes, thereby maintaining liver homeostasis and integrity. The quality control mechanism of liver regeneration likely blocks proliferation of damaged hepatocytes, preventing generation of dysfunctional or aberrant cells from the impaired hepatocytes. In this work, we have identified NIK as a hepatocyte-intrinsic sensor for liver stress and injury that controls the quality control machinery. Supporting this notion, we found that metabolic stress and numerous hepatotoxic stimuli potently activate hepatic NIK (Sheng et al., 2012). Hepatocyte-specific deletion of *NIK*, which is expected allow damaged hepatocytes to regain their proliferating capability due to disruption of their quality control mechanism, substantially increases hepatocyte proliferation and accelerates liver regeneration in *NIK^Δhep^* mice following PHx.

We have gained important insight into the potential molecular mechanism by which NIK suppresses hepatocyte proliferation and liver regeneration. NIK is known to activate IKKα (Sun, 2012). We found that mice with hepatocyte-specific deletion of *IKKα* phenocopy *NIK^Δhep^* mice with regard to reparative hepatocyte proliferation and liver regeneration following PHx. It is well established that activation of the JAK2/STAT3 drives hepatocyte proliferation and liver regeneration (Cressman et al., 1996; Riehle et al., 2008; Shi et al., 2017; Wang et al., 2011). Remarkably, we observed that IKKα binds to JAK2 and inhibits the ability of JAK2 to phosphorylate STAT3 in cell cultures. In line with these findings, phosphorylation of endogenous hepatic JAK2 and STAT3 is markedly elevated in both *IKKα^Δhep^* and *NIK^Δhep^* mice following PHx. These results unveil novel crosstalk between the NIK/IKKα pathway and the JAK2/STAT3 pathway. In light of these findings, we propose that the NIK/IKKα cascade, which is activated in damaged hepatocytes, functions as a brake to block proliferation of damaged hepatocytes through, in part, inhibiting the JAK2/STAT3 pathway. Notably, we found that NIK also directly binds to JAK2 and inhibits JAK2 activity (i.e. its autophosphorylation and ability to phosphorylate STAT3) in cell cultures. Therefore, NIK is able to suppress the JAK2/STAT3 pathway both directly and indirectly via IKKα. Additional studies are warranted to determine the relative contributions of the IKKα-dependent and the IKKα-independent mechanisms to suppression of liver regeneration by NIK.

We have provided proof of concept evidence showing that abnormally-activated hepatic NIK is responsible for suppression of liver regeneration in chronic liver disease. Chronic liver disease was modeled using two distinct approaches: chronic treatment with either hepatotoxin AAF or a HFD. In these contexts, we postulate that hepatic NIK serves as a rheostat for liver regeneration to counterbalance overgrowth of the liver. Thus, aberrant activation of hepatic NIK is expected to impair reparative hepatocyte proliferation, contributing to liver disease progression. Supporting this notion, we found that hepatocyte-specific inactivation of NIK substantially increases hepatocyte proliferation in both AAF-treated or HFD-fed *NIK^Δhep^* mice following PHx. Consistently, both phosphorylation of hepatic JAK2 and STAT3 and expression of hepatic cyclin D1 are markedly elevated in *NIK^Δhep^* mice. These findings raise an intriguing possibility that pharmacological inhibition of hepatic NIK may provide a novel therapeutic strategy to treat chronic liver disease.

In conclusion, we have identified hepatic NIK as an unrecognized hepatocyte-intrinsic sensor for hepatic stress/injury that suppresses liver regeneration through, in part, inhibiting the JAK2/STAT3 pathway. In chronic liver disease, aberrantly-activated hepatic NIK impairs liver regeneration, contributing to liver disease progression.

## Experimental Procedures

### Animals

Animal experiments were conducted following the protocols approved by the University of Michigan Institutional Animal Care and Use Committee (IACUC). Mice were housed on a 12-h light-dark cycle and fed a normal chow diet (9% fat; Lab Diet, St. Louis, MO) or a HFD (60% fat in calories; D12492, Research Diets, New Brunswick, NJ) *ad libitum* with free access to water.

### PHx models

We followed published 2/3 PHx protocols (Mitchell and Willenbring, 2008). Briefly, *NIK^flox/flox^, NIK^Δhep^, IKKα^flox/flox^*, and *IKKα^Δhep^* male mice (8 wks, C57BL/6 background) were anesthetized with isoflurane, followed by a ventral midline incision. The median and left lateral lobes (70% of the liver) were resected by pedicle ligations. Mice were euthanized 24, 48, or 96 h after PHx, and tissues were harvested for histological and biochemical analyses. Mice were introperitoneally injected, 12 h before euthanization, with BrdU (40 mg/kg body weight, ip) to label proliferating cells. A separate cohort was fed a HFD for 10 weeks prior to PHx. An additional cohort was treated with hepatotoxin 2-acetylaminofluorene (AAF) (10 mg/kg body weight, gavage) daily for 10 days prior to PHx.

Estimation of total liver weight before PHx: resected liver weight ÷ 70%. Calculation of the remnant liver weight after PHx: total liver weight - resected liver weight. Liver weight gains: terminal liver weight - the remnant liver weight. Liver growth rates: liver weight gains normalized to the remnant liver weight after PHx.

### Immunostaining

Liver frozen sections were prepared using a Leica cryostat (Leica Biosystems Nussloch GmbH, Nussloch, Germany), fixed in 4% paraformaldehyde for 30 min, blocked for 3 h with 5% normal goat serum (Life Technologies) supplemented with 1% BSA, and incubated with the indicated antibodies at 4°C overnight. The sections were incubated with Cy2 or Cy3-conjugated secondary antibodies.

### Cell cultures, transient transfection, and adenoviral transductions

Primary hepatocytes were prepared from mouse liver using type II collagenase (Worthington Biochem, Lakewood, NJ) and grown on William’s medium E (Sigma) supplemented with 2% FBS, 100 units ml^−1^ penicillin, and 100 μg ml^−1^ streptomycin, and infected with adenoviruses as described previously (Zhou et al., 2009). HEK293 cells were grown at 37°C in 5% CO_2_ in DMEM supplemented with 25 mM glucose, 100 U ml^−1^ penicillin, 100 μg ml^−1^ streptomycin, and 8% calf serum. For transient transfection, cells were split 16-20 h before transfection. Expression plasmids were mixed with polyethylenimine (Sigma, St. Louis, MO) and introduced into cells. The total amount of plasmids was maintained constant by adding empty vectors. Cells were harvested 48 h after transfection for biochemical analyses.

### Immunoprecipitation and immunoblotting

Cells or tissues were homogenized in a L-RIPA lysis buffer (50 mM Tris, pH 7.5, 1% Nonidet P-40, 150 mM NaCl, 2 mM EGTA, 1 mM Na_3_VO_4_, 100 mM NaF, 10 mM Na_4_P_2_O_7_, 1 mM benzamidine, 10 μg ml^−1^ aprotinin, 10 μg ml^−1^ leupeptin, 1 mM phenylmethylsulfonyl fluoride). Tissue samples were homogenized in lysis buffer (50 mM Tris, pH 7.5, 1% Nonidet P-40, 150 mM NaCl, 2 mM EGTA, 1 mM Na_3_VO_4_, 100 mM NaF, 10 mM Na_4_P_2_O_7_, 1 mM benzamidine, 10 μg/ml aprotinin, 10 μg/ml leupeptin; 1 mM phenylmethylsulfonyl fluoride). Proteins were separated by SDS-PAGE and immunoblotted with the indicated antibodies.

### Real-time quantitative PCR (qPCR) and ROS assays

Total RNAs were extracted using TRIzol reagents (Life technologies). Relative mRNA abundance of different genes was measured using SYBR Green PCR Master Mix (Life Technologies, 4367659). Liver lysates were mixed with a dichlorofluorescein diacetate fluorescent (DCF, Sigma, D6883) probe (5 μM) for 1 h at 37°C. DCF fluorescence was measured using a BioTek Synergy 2 Multi-Mode Microplate Reader (485 nm excitation and 527 nm emission).

### Statistical Analysis

Data were presented as means ± sem. Differences between two groups was analyzed using two-tailed Student’s t tests. P < 0.05 was considered statistically significant.

## Author Contributions

YX, AST, LR: Study concept and design; YX, AST, FW, HS, YL, MJC: acquisition of data; YX, AST, LR: drafting of the manuscript; YS, BMO, LY: critical revision of the manuscript for important intellectual content.

## Acknowledgements

We thank Drs. Lin Jiang, Liang Sheng, Chengxin Sun, and Lei Yin and Michelle Jin for assistance and discussion. This study was supported by grants DK091591, DK114220 (LR) and DK47918 (MBO) from the National Institutes of Health (NIH), fellowship #2013/07313-4 from São Paulo Research Foundation (FAPESP) (AST), and grant 81420108006 (YL) from National Natural Science Foundation of China. This work utilized the cores supported by the Michigan Diabetes Research and Training Center (NIH DK20572), the University of Michigan’s Cancer Center (NIH CA46592), the University of Michigan Nathan Shock Center (NIH P30AG013283), and the University of Michigan Gut Peptide Research Center (NIH DK34933).

## Abbreviations

PHx: hepatectomy;
HFD: high fat diet;
NIK: NF-κB-inducing Kinase;
JAK2: Janus kinase 2;
STAT3: Signal transducer and activator of transcription 3;
AAF: 2-acetylaminofluorence.

